# Temporal effects of sugar intake on fly local search and honey bee dance behaviour

**DOI:** 10.1101/2023.03.13.532281

**Authors:** Manal Shakeel, Axel Brockmann

## Abstract

Honey bees communicate navigational information of profitable food to nestmates via dance, a small scale walking pattern. Hungry flies and honey bee foragers initiate a sugar-elicited local search that involves path integration and show similarities with dance behaviour. Using a comparative approach, we explored the temporal dynamics of initiation of local search and dance in flies and honey bees, respectively. Passive displacement experiments showed that feeding and initiation of search can be spatially dissociated in both species. Sugar intake increased the probability to initiate a search but onset of walking starts the path integration system guiding the search. When prevented from walking, the motivation to begin a path integration-based search was sustained for 3 min after sugar intake in flies and bees. In flies, the behavioural parameters of search were significantly reduced for 3 min but were higher than flies that were given no sugar stimulus, indicating some degree of meander. These results suggest that sugar elicits two independent behavioural responses: path integration and increased turning, and initiation and duration of path integration system is temporally more restricted. Honey bee dance experiments demonstrated that the motivation of foragers to initiate dance was sustained for 15 min, whereas the number of circuits declined after 3 min. Based on our findings, we propose that the food-intake during foraging has the capability to activate the path integration system in flies and honey bees, and this interaction might have been elaborated during evolution to guide the walking pattern of the honey bee dance.

## Introduction

Workers of eusocial bees have evolved elaborate communication behaviours to recruit nest-mates to leave the hive and forage food for the colony. Honey bees are unique in that they are capable of communicating the location of profitable food sources to nest-mates via the waggle dance, which indicate the straight flight path to the food source (Frisch, 1967; Riley et al., 2005). The duration and direction of the waggle run phase correlate with the flight distance and direction to the food source and are presumably under the control of the brain’s path integration system which is used for navigating during the foraging trips (Riley et al., 2005; Menzel et al., 2011; Chatterjee et al., 2019). The initiation of dance behaviour, the number of waggle runs, and the intensity of the movements depend on the food quality, the arousal state of the forager and her interactions with receiver bees in the hive (Frisch, 1967, Dyer, 2002; Seeley, 1995). Recruitment behaviours of stingless bees and bumble bees show similar relations between the initiation and intensity of the locomotor displays and the food reward but do not contain navigational information (Lindauer and Kerr, 1958, 1960; Dornhaus and Chittka, 2001; Hrncir et al., 2004, 2011).

Interested in the question of how social behaviour or communication evolved from solitary behaviour, the American entomologist Vincent Dethier drew attention to similarities between a sugar-elicited local search behaviour in flies and the honey bee dance. After briefly feeding on sugar, hungry flies start a local search involving increased turning behaviour to explore the vicinity for more food (Dethier, 1957). The intensity of the search, i.e. number of turns and duration of turning behaviour, positively correlates concentration of the sugar solution and the starvation period (Nelson, 1977). Furthermore, Dethier showed that the flies started the search after they were transferred away from the location of the sugar drop, indicating that the behaviour response was not strictly dependent on the location and can be initiated spatially and temporally separated from the intake of the food. Most recently, Kim and Dickinson as well as our lab provided experimental evidence that this search behaviour also involves path integration, which means that the insects’ monitor the walking trajectory to compute distance and direction information to return to the location where they first found and ingested the food (Mittelstaedt and Mittelstaedt, 1980; Müller and Wehner, 1988; Kim and Dickinson, 2017; Brockmann et al., 2018). Murata and colleagues demonstrated that this path integration controlled search behaviour is initiated by pharyngeal and not peripheral taste receptor cells (Murata et al., 2017).

In this study, we asked two questions related to initiation of local search in *Drosophila melanogaster* and *Apis mellifera*. First, whether the location where the animal ingested the food is used as a reference point for a path integration-based local search? Second, if sugar intake and initiation of search behaviour can be dissociated, how long does the motivational effect of the food reward last? Finally, we explored in honey bees, the temporal dynamics of initiating local search and dance behaviour after sugar intake.

Using displacement experiments, we showed that in flies and bees, ingestion of sugar initiates the behaviour, in that it increases the probability to start a local search, but the path integration-based search begins with the onset of walking. By impeding locomotor behaviour post-feeding, we demonstrate that the heightened motivation to initiate search lasted for at least for 3 min in both species. A similar delay experiment with foraging honey bees showed a significant reduction in the probability to initiate dance behaviour after 15 min while the number of circuits significantly declined at 3 min.

## Results

### Location of the sugar reward had no bearing on path integration-based local search in flies and bees

To characterise path integration-based search and obtain the proportion of animals initiating search, we used a criterion of at least one return to the origin of the search and meander>0.85 for flies, and meander>0.8 for bees. In the displacement experiments, (Fig. 1A,D) 55% flies (22/40) and 56.25% bees (9/16) initiated a search involving path integration. This proportion was not different from undisplaced controls, where 61.54% (32/52) of the flies and 77.14% (27/35) of the bees initiated a search (p=0.5299 for flies; p=0.1325 for bees, Chi-square test). Hungry flies (N=11) and bees (N=12) when positioned in the arena without a sugar drop did not initiate a search (Fig. S2, S3). In addition, there were no returns and all the analysed parameters of the trajectories: meander, stay time and path length, were significantly lower compared to those of flies and bees that were provided the sugar stimulus (Fig. S4).

**Figure 1:**
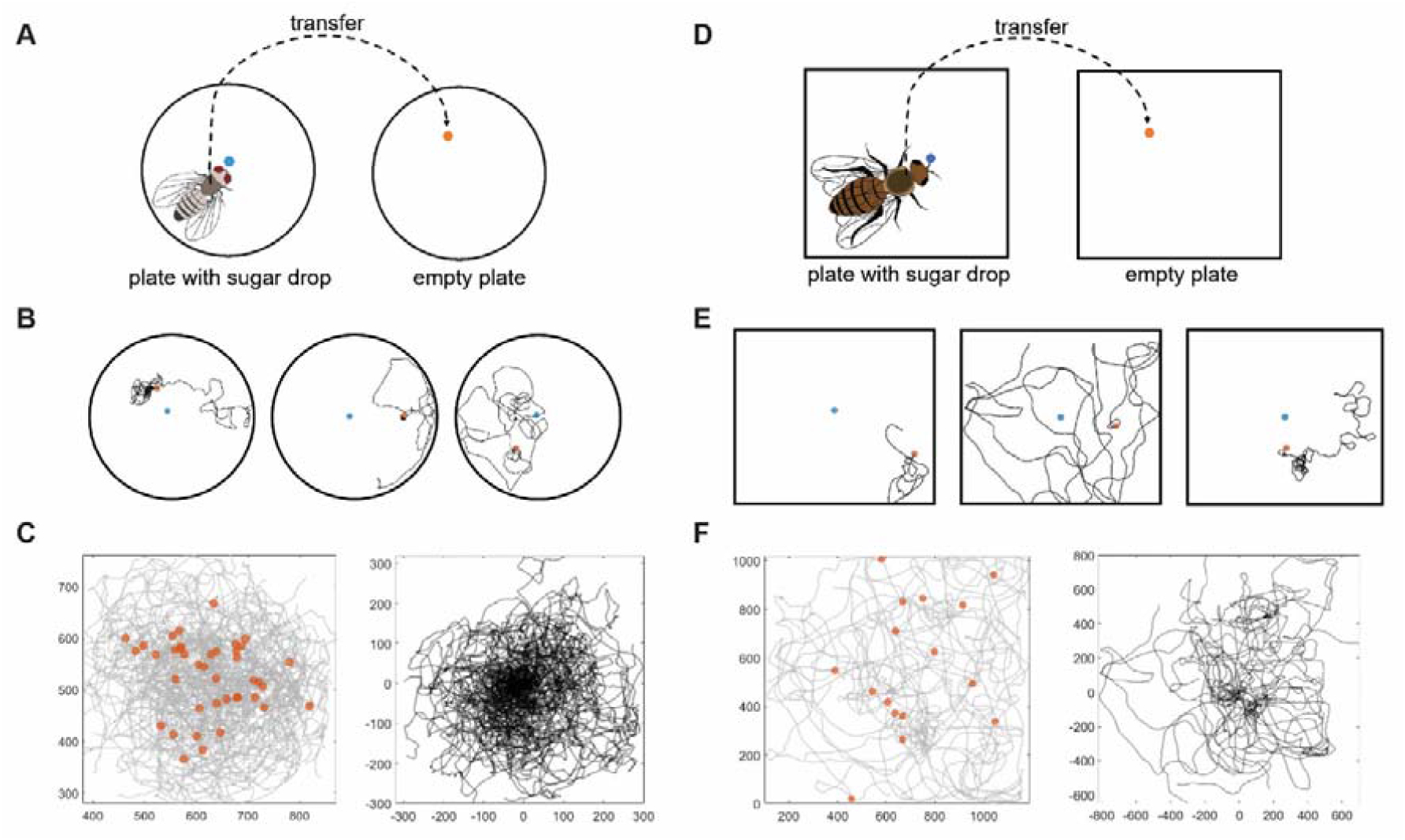
Upon passive displacement post-feeding, flies and bees start a search involving path integration around the point where they are released. (A) Schematic of transfer experiment in flies. (B) Individual trajectories showing the location of sugar drop (in blue) and position where they are released in the new arena (in orange). The x and y coordinates have been normalised to the position where each fly was transferred. (C) Left panel shows the overlay of all trajectories (N=40) without normalising to the released point. The position where each fly was set down is represented with orange circles. The right panel shows the overlay of trajectories using a new coordinate system where the origin is the position where flies were released. (D) Set up for the transfer experiment in honey bees. (E) Sample trajectories of tested bees with sugar position (in blue) and the point where they were released in the new arena (in orange). The tracks are normalised to the location where each bee was released. (F) Left: Tracks of all the tested bees (N=16) before normalising, along with the location of sugar in orange. Right: Trajectories of the bees after normalising to the point where they were released in the new arena.

Individual trajectories post-displacement (Fig. 1B,E) showed that flies and bees started a search around the position where they are transferred (orange circle) and not at the location of the sugar drop (blue circle). Overlay of the trajectories normalised for release point of all tested flies and bees confirmed these results (Fig. 1C,1F). Number of returns were similar in both species whereas meander, stay time and path length were significantly higher in flies and bees (Fig. S5).

Some of the runs of the displacement experiment allowed us to explore whether the position where the animal was set down or the position where it started to walk is used as a reference point for path integration-based search. 90% flies and 81.25% bees started walking almost immediately upon release (see methods). However, we found 4 flies (out of 40) and 3 bees (out of 16) for which we could mathematically distinguish the position where they were released and position where they started to walk. In all these cases, the returns in the trajectory were directed towards the starting position of the search (Fig. S6A,B).

### The heightened motivation to start a path integration-based local search lasted at least for 3 min post-feeding

If the sugar intake is not a releasing stimulus, the question arises for how long it affects the initiation of search behaviour. We did a set of delay experiments where flies and bees were prohibited from walking post-feeding for different durations (30 s, 1 min, 2 min and 3 min) before they were introduced into the arena. Percentage of flies and bees that initiated a path integration-based search at each tested delay duration declined linearly (Fig. 3A,B). Only 19.04% flies (4/21) and 10% bees (1/10) initiated a search when delayed for 3 min post-feeding (Fig. S7). This proportion was significantly reduced from instant transfer at this delay duration (p<0.001 for flies; p<0.05 for bees, Chi-square test). The negative slope of this decline was similar in both species (*D*.*m*.: -12.18; *A*.*m*.: -14.37, p=0.98, t-test for linear regression). Heatmaps of walking trajectories (Fig. 3C,D) showed that as the delay duration increased, the search became less centred towards the origin of walking for both flies and bees.

**Figure 2:**
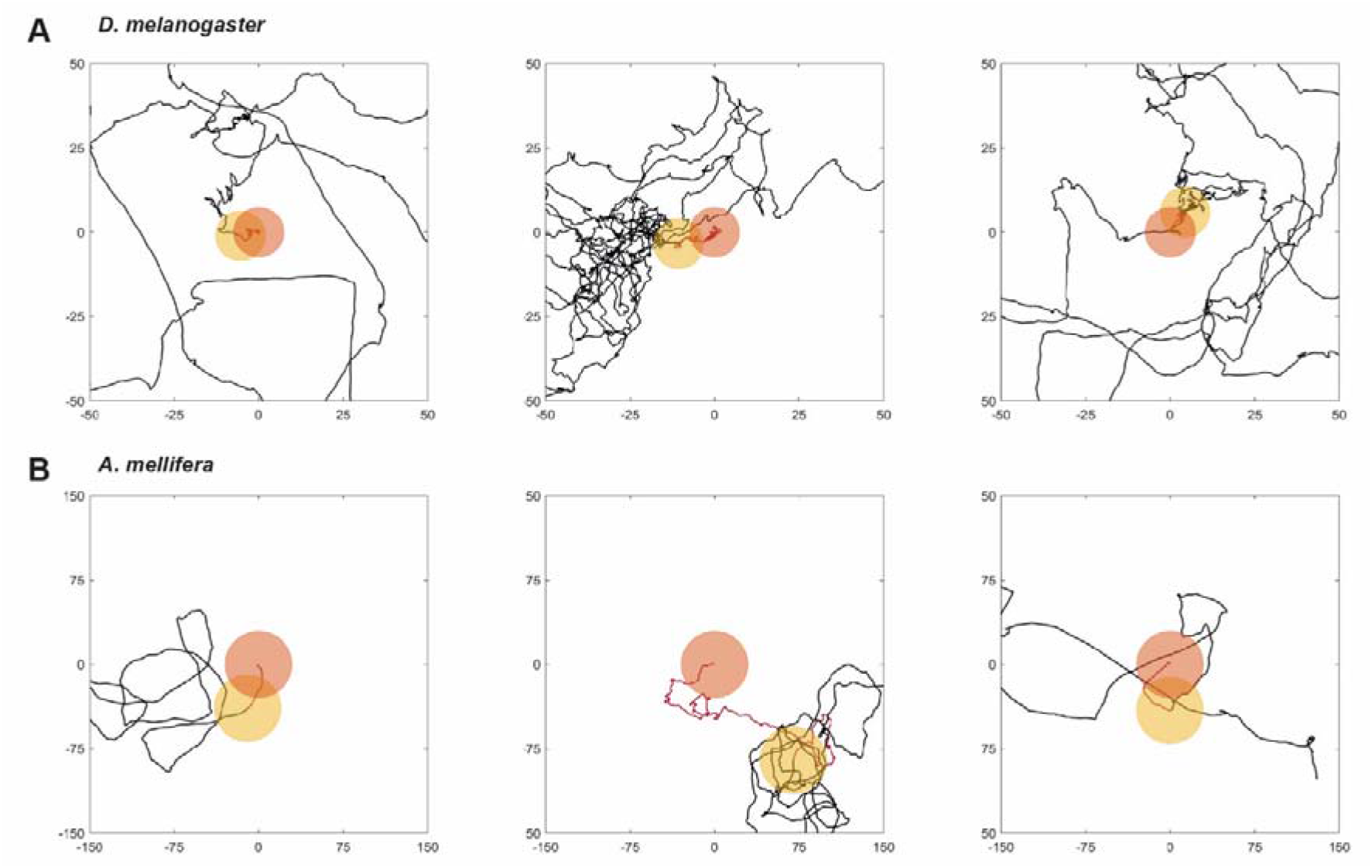
Flies and bees start a path integration-based search around the point where they begin walking. Zoomed in trajectories of (A) flies and (D) bees that walked a distance more than average time before walking ((mean±SE) 7.71±3.25 s for flies; 0.95±0.83 s for bees). The distance walked before the start of walking was 4.40±1.6 mm for flies and 7.39±6.08 mm for bees. Orange circle is drawn around the released position and the initial trajectory before walking is marked in maroon. The origin of walking is marked by a yellow circle around it.

**Figure 3:**
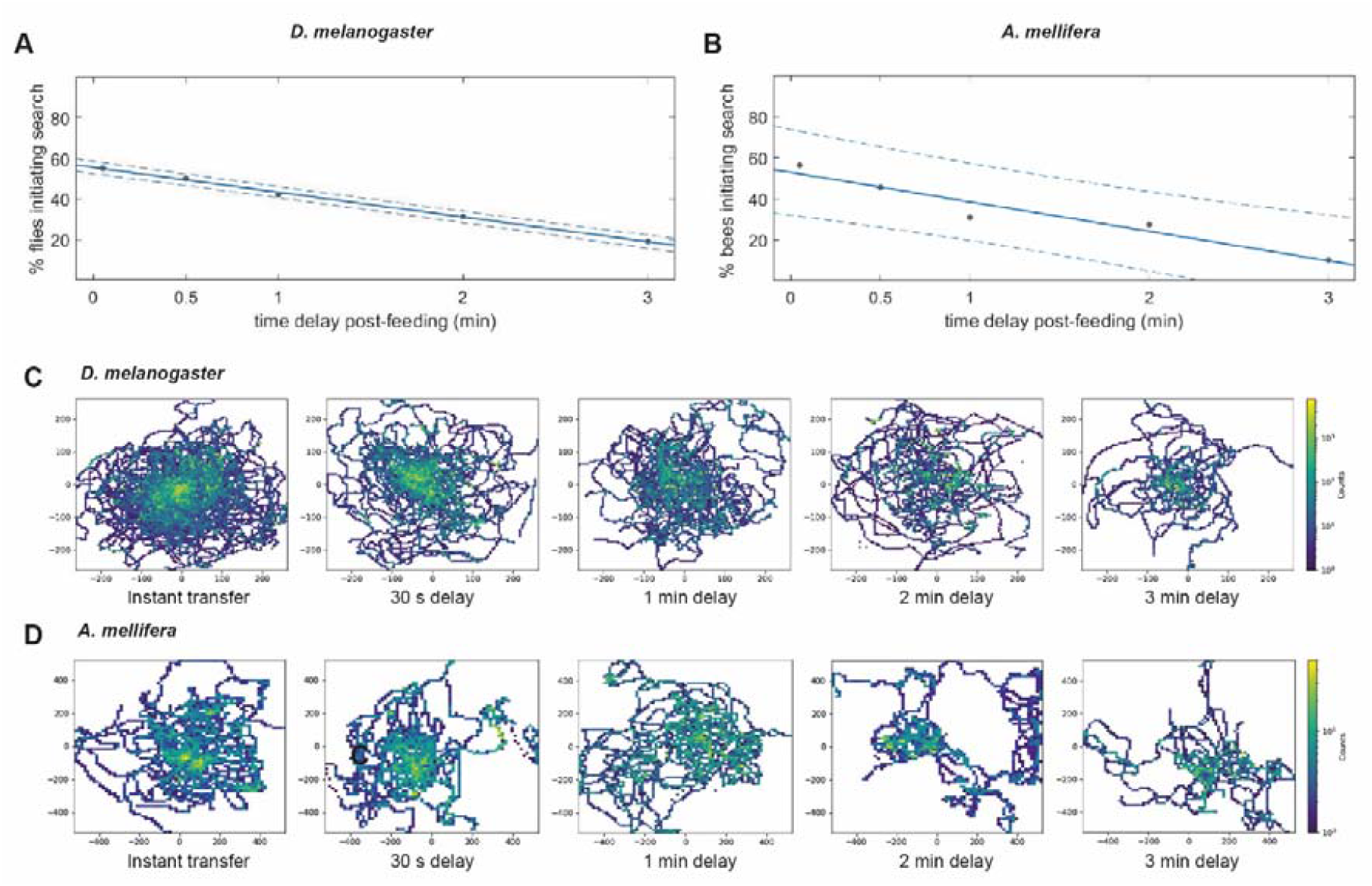
Time dynamics of initiation a path integration-based local search in flies and honey bees. (A) Percentage of flies that initiate the search decrease with the increased delay post-feeding and fits a linear model f(x) = m*x + c where m=-12.18 (−13.23, -11.14), c= 55.44 (53.67, 57.21), adjusted R-squared: 0.9971 with 95% confidence bounds, p<0.00001, t-test for linear regression. (B) The proportion of bees that initiate search decreases with increased delay durations post-feeding. The linear equation is f(x) = m*x + b2 where m=-14.37 (−21.55, -7.196), b= 52.77 (40.66, 64.89), R-squared: 0.9312 with 95% confidence bounds, p<0.001, t-test for linear regression. (C) Heatmaps of walking trajectories of all tested flies for tested durations: 30 s (N=18), 1 min (N=19), 2 min (N=16) and 3 min (N=21) show that as the time delay increases, the trajectories spread out from the origin and flies spend less time near the origin. (D) Trajectory heatmaps of bees for tested delay durations: 30 s (N=11), 1 min (N=13), 2 min (N=11) and 3 min (N=10) show that as the time delay increases, the trajectories become more spread out from the origin and bees show less occupancy near the origin.

Duration of delay had an effect on the intensity and duration of the behaviour. Number of returns were significantly reduced for both flies and bees for the 3 min delay condition (Fig. 4A-E). Meander, stay time and path length declined significantly for the three min delay period in flies, but not in bees (Fig. 4F-H). Regression analysis showed that the decline in these parameters was not strongly correlated with delay durations (Table S1). In flies, these behavioural parameters were significantly higher than unfed controls (Fig. S8).

**Figure 4:**
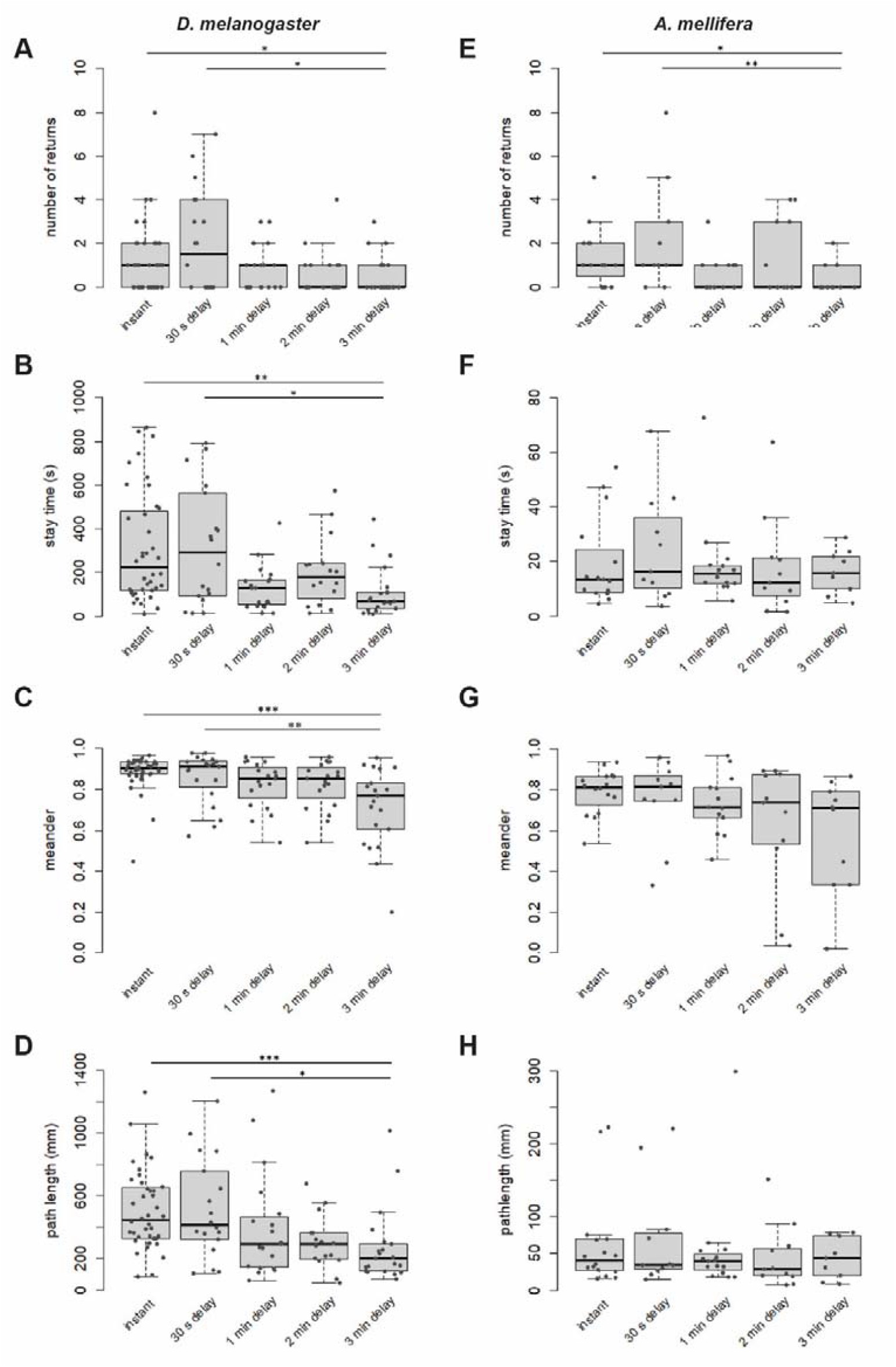
Effect of delay post-feeding on parameters of search. (A) Number of returns (B) meander (C) stay time and (D) path length for flies for 3 min delay is reduced compared to instant transfer and a short delay (30 s). (D) Number of returns in bees reduced for 3 min as compared to instant transfer and a short delay (30 s). (F-H) meander, stay time and path length, did not change with delay duration in honey bees. *p<0.05, **p<0.001, ***p<0.0001, Kruskal-Wallis test with Dunn correction, p-values adjusted with the Holm method.

### Delaying departure of foragers after sugar intake at the feeder reduced the probability of dancing

Based on Dethier’s argument that search behaviour and dance are similar with respect to the spatial and temporal separation of food intake and initiation of the behaviour, we tested the effect of delay durations between sugar collection at the feeder and return to the hive to initiate dance. Foragers were assigned into two groups: treatment and control, based on their dance activity. We calculated the probability of dancing post-delay as the ratio of the number of foraging trips with dances post-treatment to the number of total trips for the respective delay treatment. The probability of dancing reduced linearly with delay duration (Fig. 5A) but was not significant lower at 3 min (p=0.9656). We observed a significant reduction in the probability after a delay of 15 min (dance probability=0.4, p<0.05 instant-15 min, p<0.001 30 s-15 min, Chi-square test).

**Figure 5:**
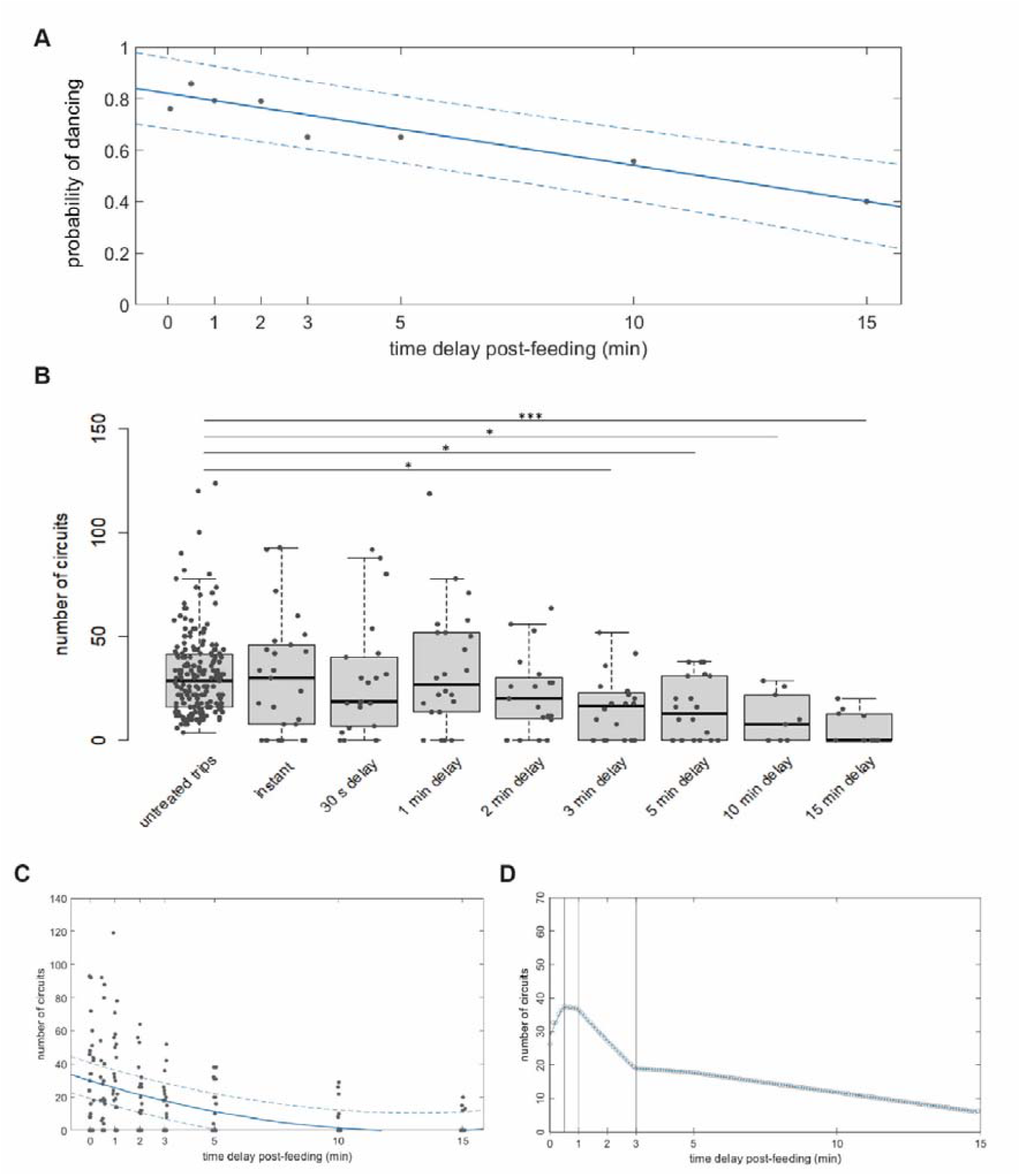
Time dynamics of initiation of dance in honey bees. (A) The probability of dancing for bees decreased with delay duration with a linear model: f(x) = m*x + b where m= -0.1485 (−19.5, -10.2), b=0.6918 (64.83, 73.54), R-squared: 0.8878 with 95% confidence bounds; p<0.00001, t-test for linear regression. (B) Number of circuits showed a significant reduction when the delay increased to 3 min and reduced further as the delay was increased to 15 min. *p<0.05, ***p<0.0001, Kruskal-Wallis test with Dunn correction, p-values adjusted with the Holm method. (C) Decline in the number of circuits decreases with time delay post-feeding. Linear model Poly2: f(x) = a1*x^2 + c2*x + c3 where a=2.855 (2.02, 3.69), b=-14.16 (−15.9, -12.41), c=17.09 (15.87, 18.3), R-squared: 0.9491 with 95% confidence bounds. (D) 3 min is detected as one of the change-points with a p=0.98042. Two other change points were identified as 30 s and 1 min (p=0.99950 and p=0.99946 respectively).

There was no change in the number of circuits between the dances of the control bee and the dances of untreated trips of the experimental bees (Fig. S9). Therefore, we compared the number of circuits of treated foraging trips with the untreated trips to estimate the effect of delay. We observed a significant decline in the number of circuits for the delay period of 3 min (Fig. 5B). Longer delay durations (5 min, 10 min and 15 min) showed a further decrease in circuits with respect to untreated trips, with the strongest reduction after 15 min (Fig. 5B, Table S2). Regression analysis showed a strong correlation between delay duration and decline in the number of circuits (Fig. 5C, Fig. S10). Change-point analysis detected a pronounced reduction in the number of circuits for delay durations of 1 min and 3 min (Fig. 5D).

## Discussion

Passive displacement experiments have been successfully used to identify navigational mechanisms underlying food-seeking and foraging behaviours (Müller and Wehner, 1988, 1994; Dacke et al., 2020; Patel et al. 2022). We ourselves recently used displacement experiments to demonstrate that the sugar-elicited search behaviour involves path integration to guide the animal back to the location of where they had previously found food (Brockmann et al., 2018). In this study, we provide experimental evidence that the onset of walking starts the path integrator. The location where the flies and bees started walking was used as the origin of search instead of the location where they found the food. In a few of our experiments, the displaced flies (4/40) and bees (3/16) made small movements away from the original location before starting the search. In those cases, the returns were distinctly oriented to the position where they started their walking trajectory. Evidently, the path integration system should be functionally coupled to the initiation of (self-) locomotion (Wehner and Srinivasan, 2003; Seidl, 2006). We propose that sugar intake releases the behaviour sequence modulating the probability to start a local search involving path integration (Dethier, 1957; Heiligenberg, 1977; Brockmann et al., 2018).

Earlier studies on the sugar-elicited search behaviour in *Drosophila* highlighted that the intensity and duration of the search depends on the starvation period, sugar concentration, the volume taken in, and the “central excitatory state” of the fly, supporting the idea that the sugar intake modulates the behaviour (McGuire and Tully,1986; Mayor et al., 1987). In our delay experiments, the modulatory effect of the sugar stimulus (*D*.*m*.: 0.2 ul, 500mM; *A*.*m*.: 3ul, 2M) to initiate the search lasted for 3 min in flies and honey bees. After this duration, we observed a significant decrease in the percentage of animals starting a path integration-based search. In addition, we observed a reduction in the number of returns for both species, while only flies showed a significant decline in meander, stay time and path length for this duration. Stay time and path lengths for honey bee trajectories were considerably shorter than those for fly trajectories (Fig. S8). It might be possible that these parameters are not sensitive enough to detect an effect of delay duration on behaviour in honey bees.

Although our analyses indicated that for the majority of flies, the 3 min delay inhibited the initiation of path integration-based search, the walking trajectories showed higher meander, stay time and path length compared to unfed controls. This finding may suggest that sugar-elicited search behaviour is composed of two independent behavioural responses, path integration guided walking and increased turning frequency, regulated by different modulatory dynamics. High turning behaviour is a common strategy for optimising search. Since it is less energy-intensive, its initiation and duration might be less temporally restricted than path integration (Bell, 1999; Sterling and Laughlin, 2015).

The honey bee dance experiments delaying the departure of foragers from an artificial sugar feeder showed that the motivation to initiate a dance in the hive lasted at least for 15 min. The differences in the effective delay durations between search and dance are certainly a consequence of the different behavioural contexts. The motivation to recruit nestmates after food collection should last longer than searching for food around a site where they found a small amount of sugar. Although the experimental conditions were different, the slopes of decline in initiation of search (−14.04) and dance (−0.1485) in honey bees was comparable (p=0.8730, t-test for linear regression), which might suggest that both behaviours are regulated by the same neuromodulatory system.

Several scenarios have been proposed on how dance communication might have evolved (Price and Grüter, 2015; Barron and Plath, 2017). Successful foragers of the closely related stingless bees and bumble bees perform excited sugar-dependent locomotor displays or runs over the nest area that alert their nestmates (Lindauer and Kerr, 1958; 1960; Dornhaus and Chittka, 2001; Menzel, 2019). However, none of these behaviours show any features that are correlated to the flight direction towards the food source. The question remains, how flight navigational information got incorporated in these excited behaviours during evolution. The finding that sugar-elicited search behaviour activates the path integration system at least suggests an ancestral connection between neural pathways of food reward and the path integration system which was recently shown to involve the central complex (Dyer et al., 2002; Heinze and Homberg, 2007; Heinze et al., 2018, Collet, 2019). Flies and bees might exhibit similar (ancestral) modulatory circuits regulating the activity of the navigational system (Barron et al., 2007; Schröter et al., 2007; Busch et al., 2009; Menzel, 1999, 2009; Yang et al., 2015). A study of the sugar-induced search behaviour with a focus on the initiation of the path integrator in *Drosophila* would be interesting in itself but also promises to help searching for the neural pathways involved in dance communication in honey bees.

## Method Details

### Fly rearing

Male flies of *Drosophila melanogaster* Canton-S (CS) strain were used in all the experiments. Both male and female flies show sugar-elicited search behaviour but starvation time is more consistent among male flies. Additionally, female flies change their feeding preference after mating (Carvalho et al., 2006). Hence, we used only male flies. Flies were reared and maintained on standard fly media at 25°C in a 75% relative humidity in a 12-hour light/dark cycle. Flies eclosing within a 12-hour period were collected and maintained in fresh medium for two days. The flies were starved of food (with access to water) before the behavioural experiments. To standardise the hunger state across trials, flies were starved for the duration of 90% survival of the population under food-starved conditions. Food-starvation tolerance was calculated by depriving two day old flies of food with access to water with soaked Kimwipe paper at the bottom. 15 flies were placed in a vial (N=3). Number of surviving flies were counted every hour and survival curves were plotted. The duration at which 90% of the starved flies survived was used as the starvation period. This starvation period for CS flies ranged from 26-29 hours in tested flies, over the course of the experiment. Behavioural experiments were done between 1400-1700 hrs.

### Experimental procedure

Flies were individually tested for local search after starvation. 90 mm Petri dishes were used as an arena for behavioural assays. The arena was illuminated from the bottom by a panel of surface-mounted cool white LEDs. The light intensity was 320 lux at the centre of the arena, measured using TENMARS TM-203 Data Logging Light Meter. 2-D position of the flies was recorded with an overhead camera at 40 fps (Flea3, Point Grey, 1214 mm lens, Azure). The Petri dish was surrounded by a white cylinder (51.5 mm height, 114 mm inner diameter) made of polyvinyl chloride resin to contain the visual field of the flies (Fig. S1A). Petri dishes were wiped with 70% ethanol between trials.

A drop of sucrose solution (0.2 μl, 500 mM) was positioned in the centre of the arena. A 2 ml microcentrifuge tube (inner diameter 8.7 mm, length reduced to 5 mm by stuffing cotton at the bottom) housing a single fly was inverted over the sugar drop. The experimenter waited until the fly found the sugar and started feeding before removing the tube. We used this protocol to record the behaviour of control flies, who were fed with sugar and allowed to search without any displacement. These flies were used as undisplaced control and control against unfed flies, who were not given any sugar. We observed the trial till the time the fly reached the periphery, climbed over the edge and left the arena. For passive displacement experiments, the fly was allowed to feed on a drop of sugar for 40 s. This time was fixed based on the mean feeding time of fed flies in control experiments. The fly was then gently picked with a mouth aspirator (diameter<0.5 mm) contacting the thorax and we removed the Petri dish. The fly was then transferred to an arbitrary position on a fresh Petri dish and allowed to walk freely. This transfer procedure took 3-4 seconds and was referred to as instant transfer (N=40). For delay experiments, the fly was allowed to feed for 40 s and then housed inside a tube (inner diameter 4.6 mm∼1.5 fly length) to impede walking. The tube was cushioned on both ends, giving little to no movement for the fly, since we did not want the fly to start walking while it was inside the tube. This prohibited the fly from initiating search for specific durations. The delay durations tested were 30 s (N=18), 1 min (N=19), 2 min (N=16) and 3 min (N=21). After the experimental delay duration elapsed, the fly was introduced to a fresh Petri dish at an arbitrary position and its behaviour was recorded. Each experimental day consisted of tests with several time delays with the starved flies from the same batch in order to reduce the effect of day and observer bias.

### Honey Bee Local search

*Apis mellifera* colonies (N=4) were obtained from a local beekeeper. The colonies consisted of 4-frames containing approximately 2000 individuals per frame. The colonies were housed on a lawn inside the campus of the National Centre for Biological Sciences, Bangalore, India. Foragers were trained to an unscented feeder containing 1 M sucrose kept 5 m from the hive entrance. The colonies were also given pollen on an artificial feeder but the pollen feeder was removed during the experimental hours. The bees were trained for two hours from either 1-3 PM or 2-4 PM depending on the weather conditions.

Nectar foragers arriving at the feeder were individually tested for local search. A single bee was collected in a tube (inner diameter: 14.9 mm) before it started collecting sugar at the feeder. It took us under a minute (an average walking time 55.27 s was recorded over days) to reach the behaviour room from the feeder location. We used a larger square arena (30.2 cm x 30.2 cm) for honey bee search experiments. The arena made of transparent acrylic sheet was backlit by a panel of surface mounted cool white LEDs. The tube housing the honey bee was placed on the illuminated arena for 1 min so that it could get acclimated to the change in lighting conditions from outside. A drop of sucrose solution (3 μl, 2 M) was placed in the centre of the arena. The bee was quickly transferred to a bigger tube (inner diameter: 27 mm, length reduced to 8 mm by stuffing cotton at the bottom). This bigger tube was then inverted over the sugar drop and the experimenter removed the tube after the bee found the sugar and started feeding. We used this method to record the search of control fed bees while no sugar was provided for unfed controls. We observed the trial till the bee flew away or reached the periphery of the arena, climbed over the edge and left. The light intensity was 520 lux as measured by TENMARS TM-203 Data Logging Light Meter, at the centre of the arena. Search was recorded with an overhead camera at 40 fps (Flea3, Point Grey, 1214 mm lens, Azure). The floor of the arena was wiped with 70% ethanol between trials.

For passive displacement experiments, we provided the sugar in a 90 mm Petri dish placed in the centre of the arena. This helped us save time since we could quickly remove the Petri Dish and introduce the bee in the arena. The bee was first allowed to feed on a drop of sugar for 8 s. This time was fixed based on the average feeding time of the fed group of control bees. The bee was gently picked with a mouth aspirator (diameter ∼2.5 mm) contacting the thorax and the Petri dish was removed. The bee was then transferred to an arbitrary position in the arena and allowed to walk freely. This transfer procedure took about 4 s and was called instant transfer (N=16). For delay experiments, we modified the protocol from flies experiments, since bees got agitated when we used the same method. Instead of housing the bees inside a tube to impede locomotion, an inverted tube (inner diameter: 14.9 mm, ∼1 bee length) was placed over the bee to immobilise it. The inverted tube with the bee was gently moved away from the sugar drop and closer to the edge of the Petri dish for the elapsed delay duration. The Petri dish was also moved in some cases to an arbitrary location in the square arena. The tube was cushioned, and there was no space for the housed bee to walk. The delay durations tested were 30 s (N=11), 1 min (N=13), 2 min (N=11) and 3 min (N=10). After the experimental delay duration elapsed, we gently slid the bee from the Petri dish to the arena underneath and simultaneously removed the tube from the top. Thus the bee was introduced to the arena and we recorded its behaviour. We tested several time delays on the same day in an arbitrary fashion to reduce the effect of day and observer bias.

### Honey Bee Dance

*Apis mellifera* colonies (N=3) were obtained from a local beekeeper. The colonies were housed in an outdoor flight cage (16 m × 4 m × 4 m) on the campus of the National Centre for Biological Sciences, Bangalore, India. A pollen feeder was advertised outside of experimental hours. The day-night length as well as the temperature conditions inside the flight cage were the same as the natural conditions in Bangalore.

The colony was transferred to an observation hive for recording the dances. We trained the foragers to an unscented sucrose solution (1M) from 11-12:30 PM using a gravity feeder kept on a channel plate (Fig S1B) 11 m away from the hive entrance. 15-20 bees were individually marked (Posca 5M, ‘Uni’ Mitsubishi Pencil, India) to form a foraging group. The experiment was done over four days and three observers were present at the feeder throughout the experiment. One person recorded the number and timing of the foraging trips. The other caught all recruits (non-marked foragers) coming to the feeder to keep the foragers motivated to dance throughout the experiment (von Frisch 1967). These bees were released back to the hive after the observation time. The third person performed the time delay procedure after the foragers collected sugar water. On day 1, we recorded the dance activity of the marked foragers. The video was then analysed to identify the most active dancers (4-6) from the group (George and Brockmann, 2019).

This was done based on ranking the bees based on the dance intensity (the ratio of number of circuits to the number of dances) and dance probability (the ratio of the number of dances to the number of foraging trips). These active dancers were randomly assigned to either the experimental or control group. On days 2-4, the experimental group underwent treatments while the control bees foraged freely. The foraging trips of the experimental group consisted of treated trips (where the delay was performed) and untreated trips (where no delay procedure was performed). We used one foraging group from colonies 1 and 3, while colony 2 was used for two foraging groups.

To introduce time delay post-feeding, we took advantage of the natural feeding behaviour of bees. After the foragers collect sugar water, they retreat their proboscis and walk a couple of steps away from the sugar solution before flying. Just as the bees were about to take flight, a tube was gently inverted over the bee, similar to how we restrained the bees post-feeding in search experiments. This tube was painted black from outside and cushioned so that there was no space for the housed bee to walk inside (14.9 mm, ∼1 bee length). The treatment was started after the bees in the experimental group completed five foraging trips. Time delays that we tested in this experiment were: 30 s, 1 min, 2 min, 3 min, 5 min, 10 min and 15 min. Since our camera allowed real-time monitoring, we decided to test longer time delays because we did not see an extinction of dance for up to 3 min. An instant treatment was also done, where the tube was inverted over the bee post-feeding for 3-4 s (same duration as instant transfer in Honey bee search experiments). After the delay duration elapsed, the tube was lifted and the forager flew back to the hive. We tested each bee in the treatment group for all the time delays on each day to eliminate the effect of the day. Trips where a forager started feeding again after we performed the delay treatment were removed from the analysis. In total, 148 treatments over 11 individuals were tested (instant: N=26, 30 s: N=21, 1 min: N=24, 2 min: N=19, 3 min: N=20, 5 min: N=19, 10 min: N=9, 15 min: N=10).

### Analysis of trajectories for local search

Ctrax -The Caltech Multiple Walking Fly Tracker (Branson et al., 2009; http://ctrax.sourceforge.net/) was used for tracking the flies. MATLAB 2019b, Rstudio (Version 1.4.1106) and Python are used for analysis.

For control experiments, we defined the end of feeding and start of walking as flies moving at a speed>4 mm s-1 and bees moving at a speed>3 mm s-1 in three consecutive frames. We used the same criterion to define walking in the transfer experiment. We calculated the time it took the insect to start walking after being transferred to the new plate as ‘latency in walking’. The distance covered in this time by flies and bees was called ‘distance before walking’.

Path length (distance walked during the search, in mm) and stay time (time spent in search, in s) were calculated from search trajectory. Meander was calculated by dividing the beeline of the path to the path length, and subtracting from 1. High values of meander indicate more tortuosity in the walking path. We developed an algorithm to identify and count the number of returns using two concentric circles. An inner circle indicating the origin of search (Rin = 2.5 mm, 10 mm for bees) and the outer circle indicating the minimum distance (Rout = 4 mm for flies, 16 mm for bees) that the animal had to move away from the origin. A return was defined as a movement out of the outer circle (Rout) and then coming back into the inner circle (Rout).

### Dance recording and analyses

We used a mounted video camera (GoPro Hero 8 Black CHDHX-801) to record the dances at 60 fps. We used this camera in order to monitor the videos in real time. We made a temporary barrier at the nest entrance between the two faces of the observation hive to direct the forager traffic to the side facing the camera. Since the food source was very close to the hive (11 m) the waggle run duration for the dances was very short (Gardner et al. 2008; Seeley 1994). Therefore, we counted the number of waggle dance circuits in each forager’s dance (Seeley, 1995, von Frisch, 1967). Videos were analysed manually using Virtual Dub 1.10.4 (http://www.virtualdub.org/). Change-point analysis was done using BEAST (https://github.com/zhaokg/Rbeast) is a Bayesian algorithm to detect change points and decompose time series into trend, seasonality, and abrupt changes (Zhao et al., 2019).

## Supplemental Data

**Figure S1:**
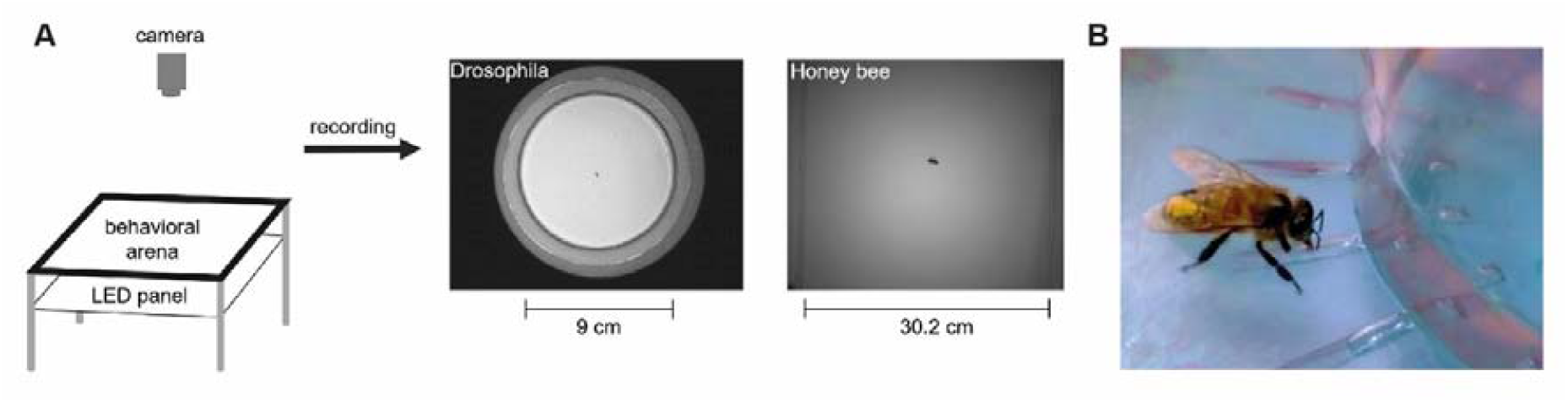
Local search and dance experiment. (A) Experimental set-up for search experiments: The arena is backlit by a panel of white LEDs. The overhead camera is connected to the computer for tracking the walking flies and bees. (B) A marked forager feeding from the channel plate before being captured for delay in dance experiments.

**Figure S2:**
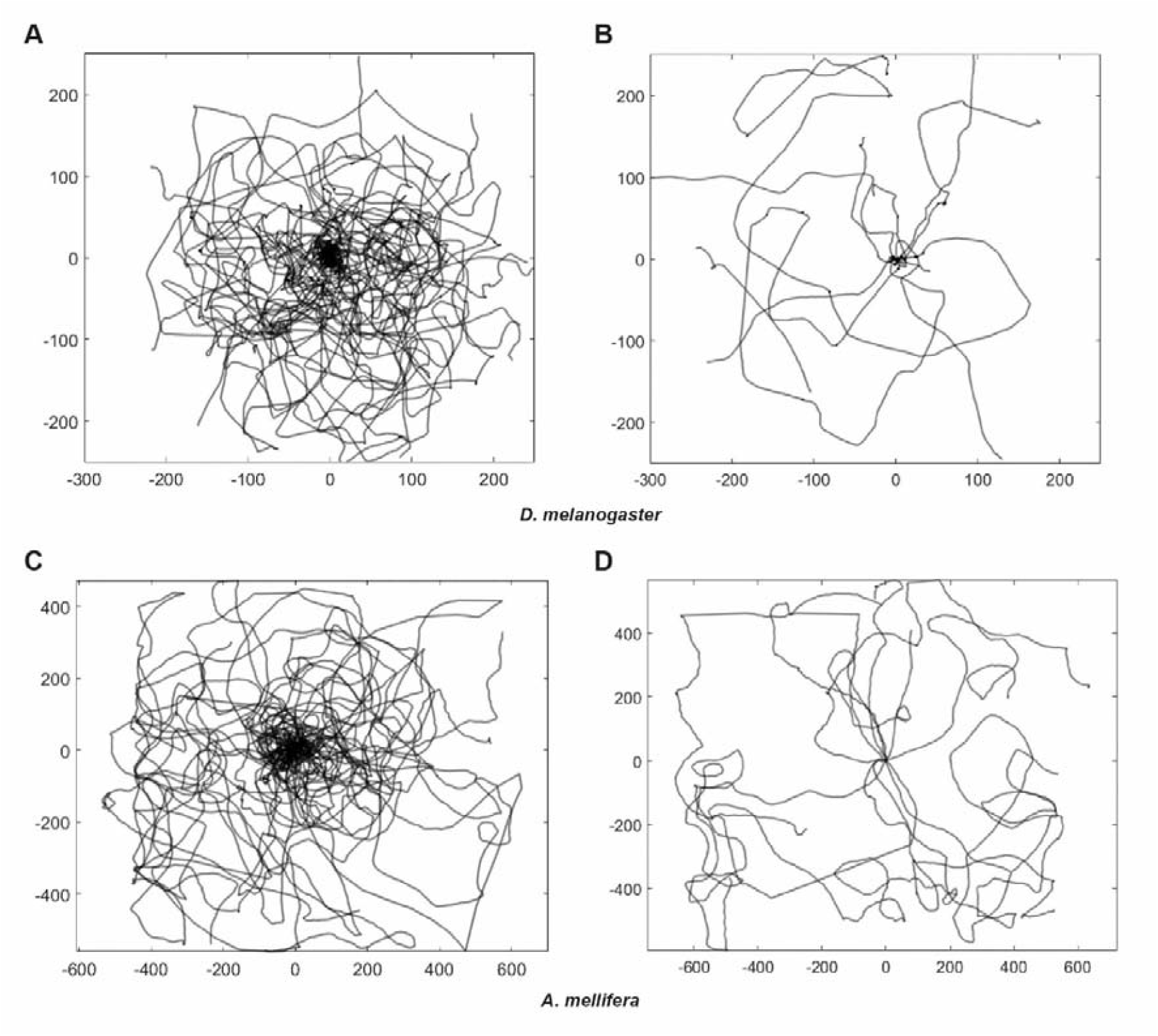
Sugar intake is necessary for initiating local search. (A) Overlay of search trajectories of 11 flies from the control group for local search stimulated with 500 mM, 0.2 μl sugar solution (B) Overlay of the paths of the flies (N=11) which were given no sugar. (C) Overlay of search trajectories of 12 bees from control fed with 2 M, 3 μl sugar solution (D) B) Overlay of path of the bees (N=12) recorded without sugar stimulation.

**Figure S3:**
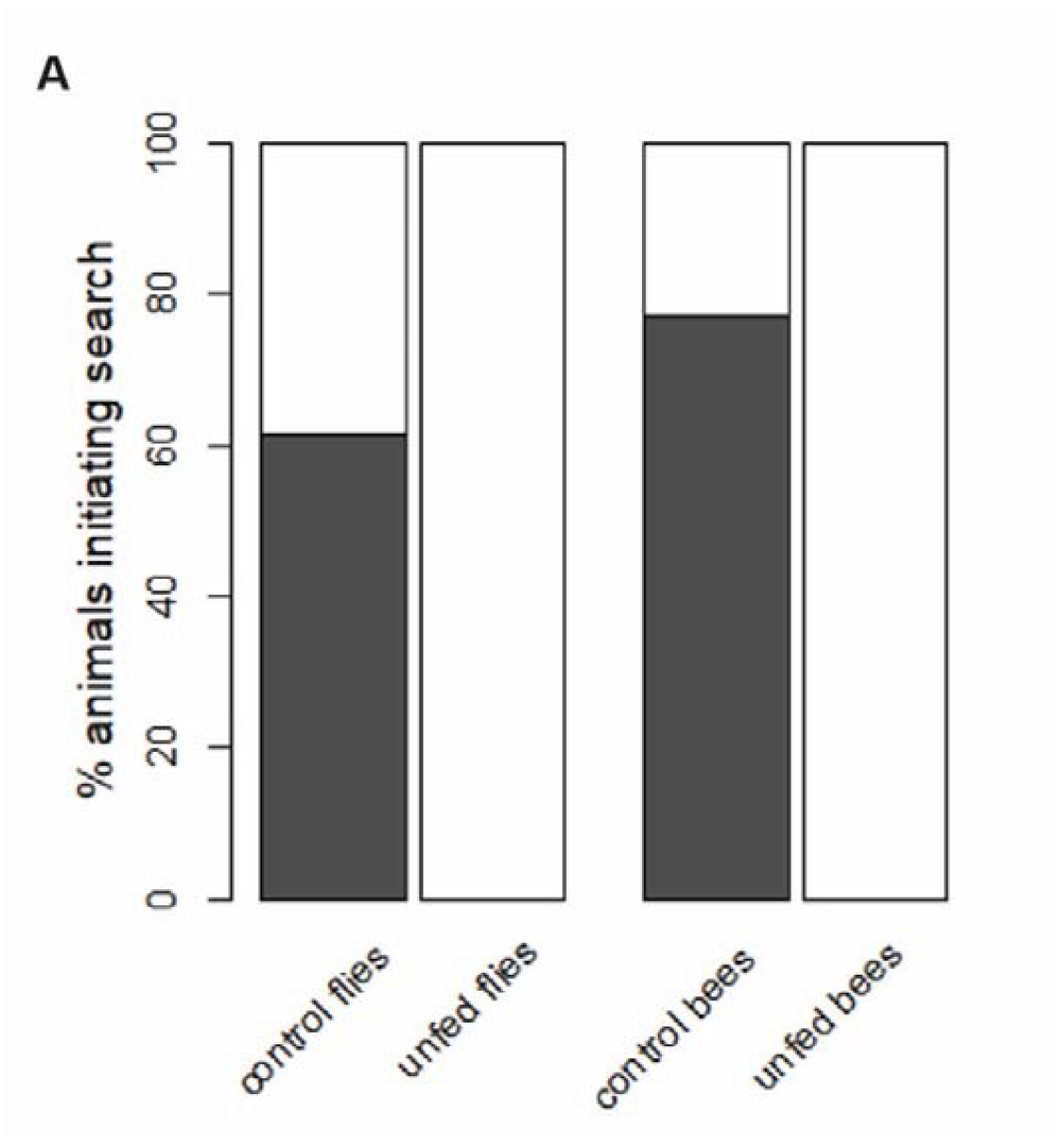
Hungry flies do not initiate search when there is no sugar stimulus. 61.54% (32/52) flies and 77.14% (27/35) bees initiated search when they were fed with sugar. 0% flies and bees initiated search in the absence of sugar reward.

**Figure S4:**
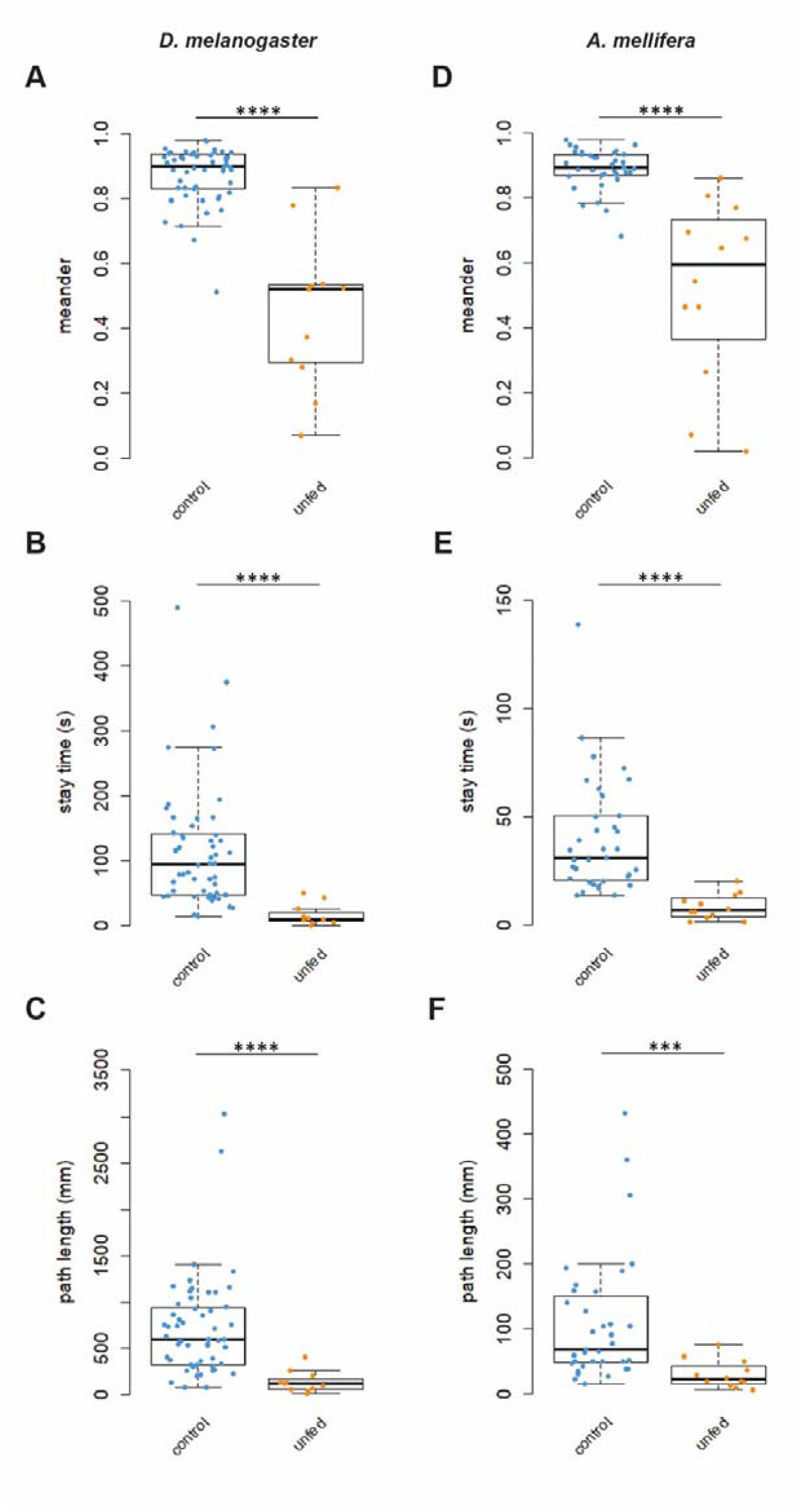
Behavioural parameters are significantly lower in unfed flies and bees. (A-F) Meander, stay time and path length were smaller for hungry flies that were not given sugar reward compared to control flies and bees that were given stimulated with sugar. ***p<0.0001, ****p<0.00001, Wilcoxon Rank Sum Test.

**Figure S5:**
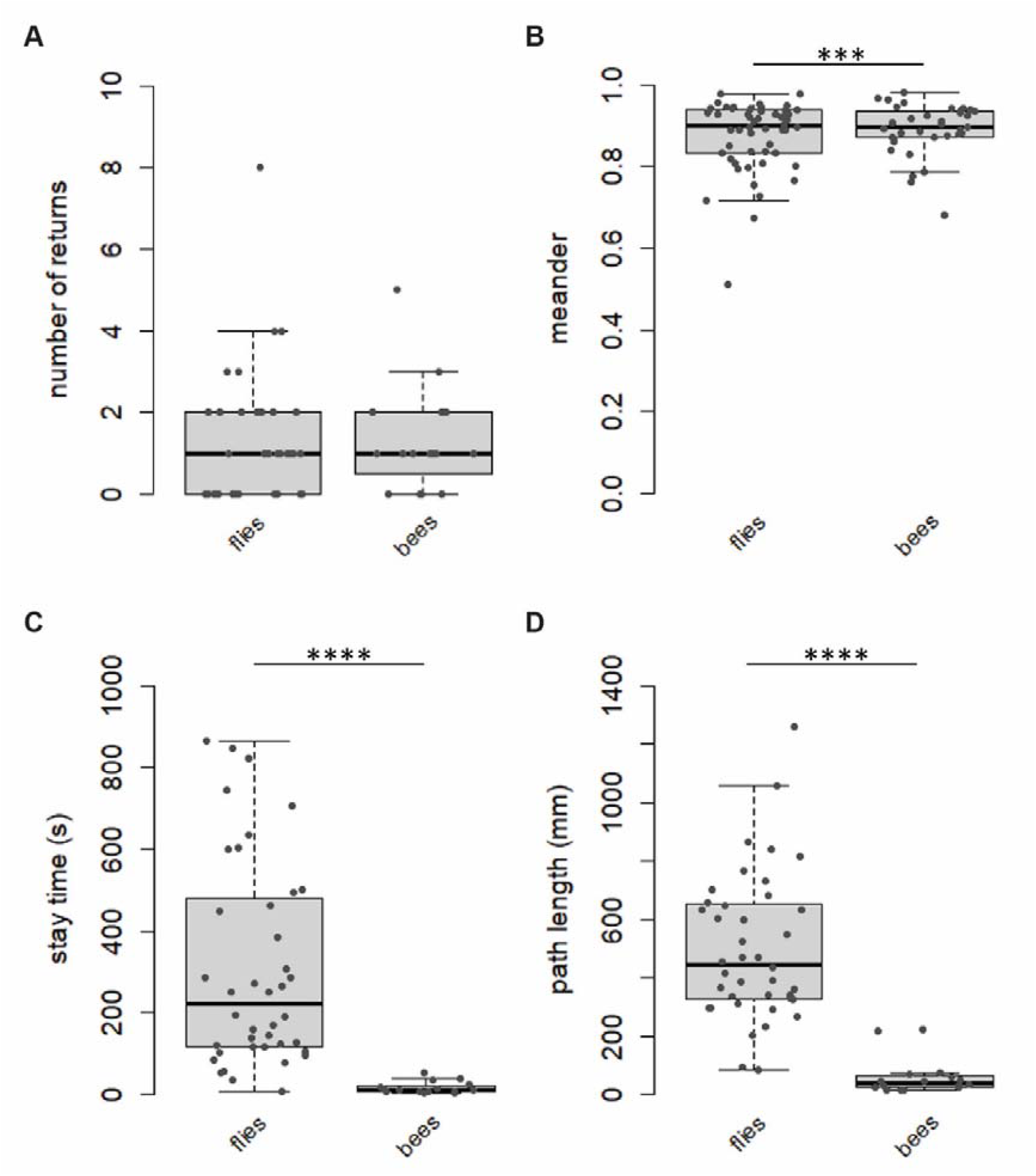
Comparison of behavioural parameters between flies and bees for displacement experiments. (A) Number of returns do not show a difference between the two species. (B-D) Meander, stay time and path length are lower in bees than flies. ***p<0.0001, ****p<0.00001, Wilcoxon Rank Sum Test),

**Figure S6:**
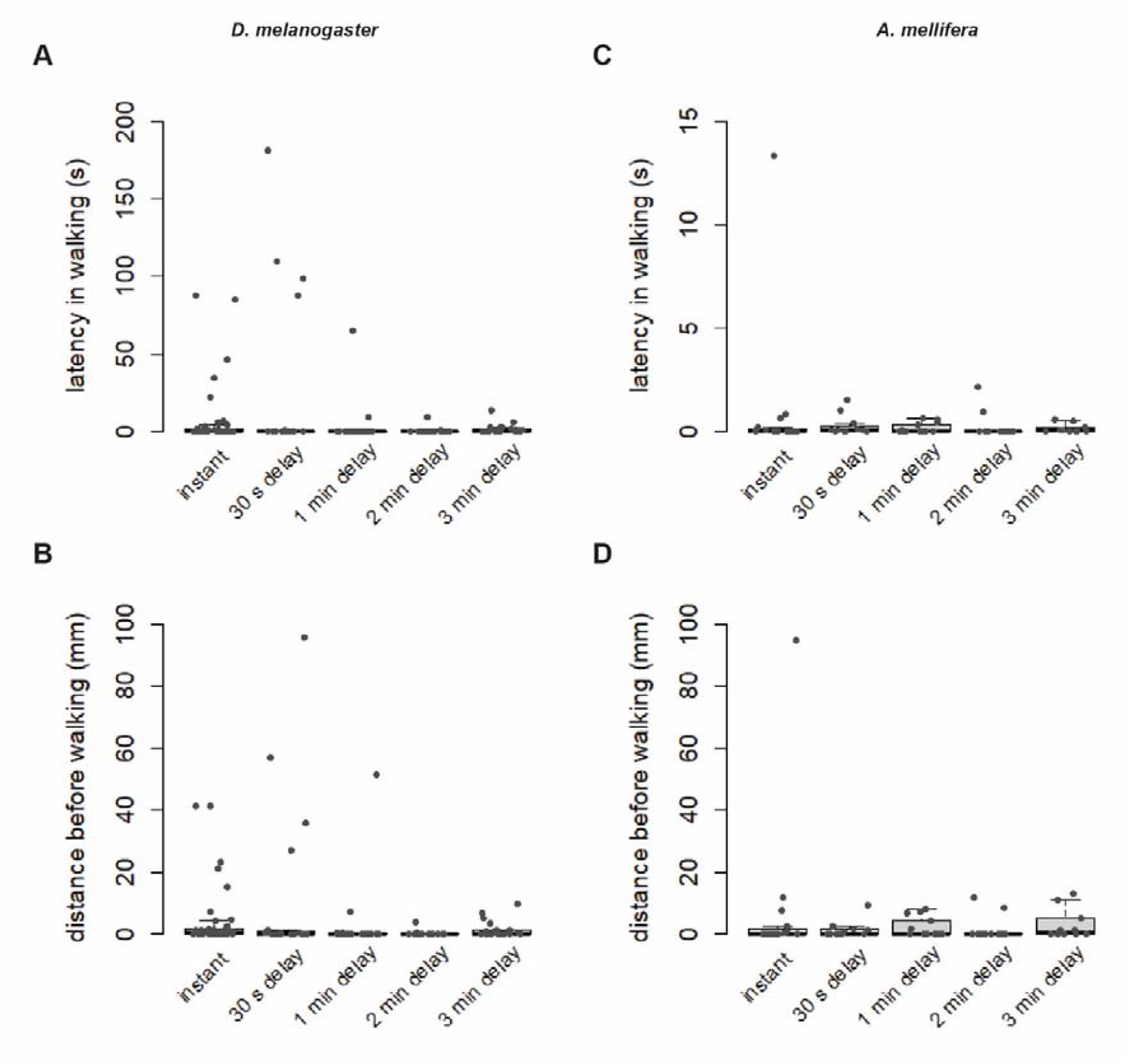
There is no difference in latency to walk post-transfer across delay durations. (A,B) Latency in walking and distance covered before walking does not change with increased time delay post-feeding in flies. (C,D) Bees do not show a difference in delay in walking and distance covered before walking with increased delay post-feeding. Kruskal-Wallis test with Dunn correction, p-values adjusted with the Holm method.

**Figure S7:**
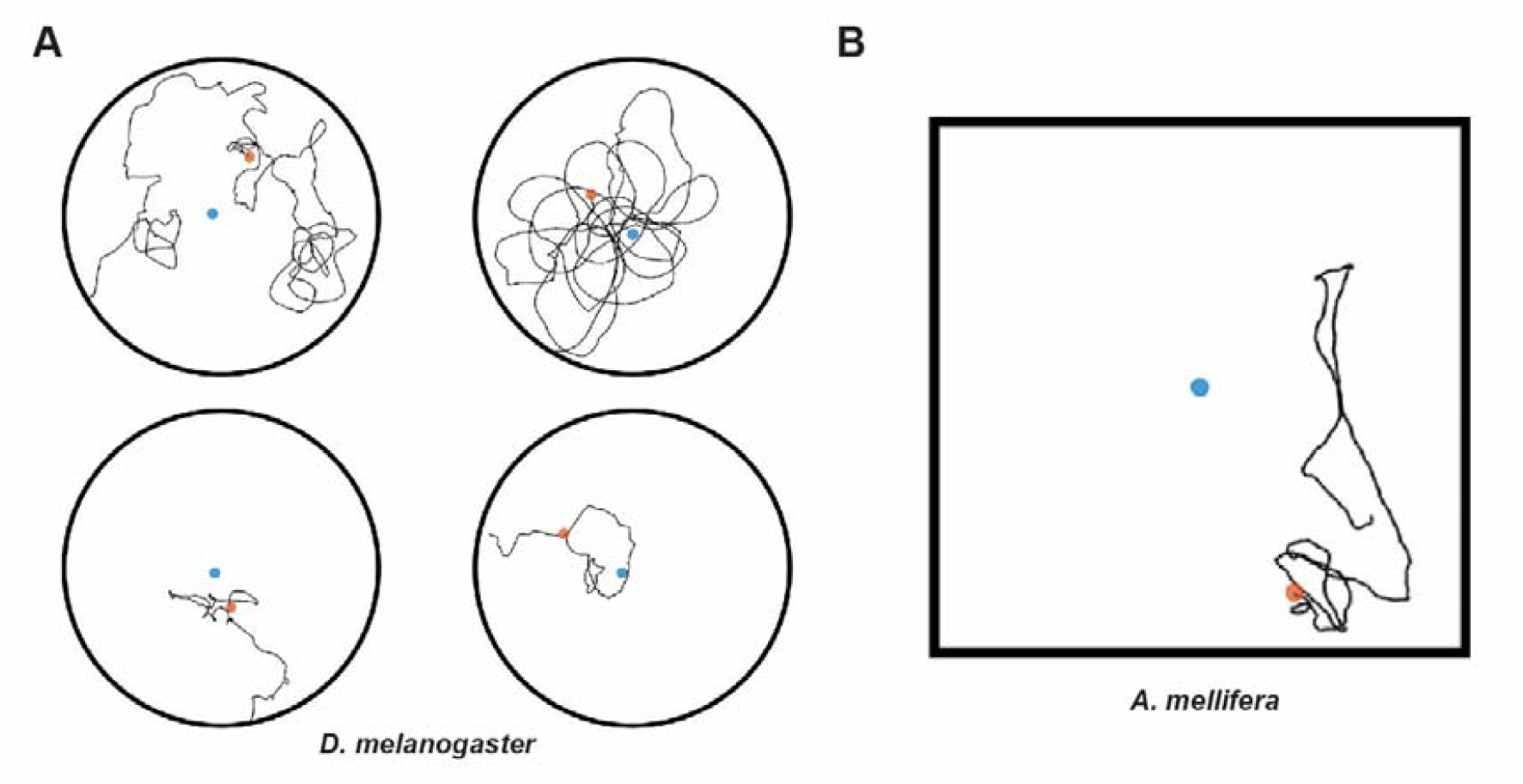
Less than 20% of tested flies and bees initiate local search for a delay of 3 min. (A) Trajectories of the four flies out of 21 tested flies for 3 min delay that initiated a path integration-based search. (B) Trajectory of the only bee (out of 10) which initiated a path integration-based search.

**Figure S8:**
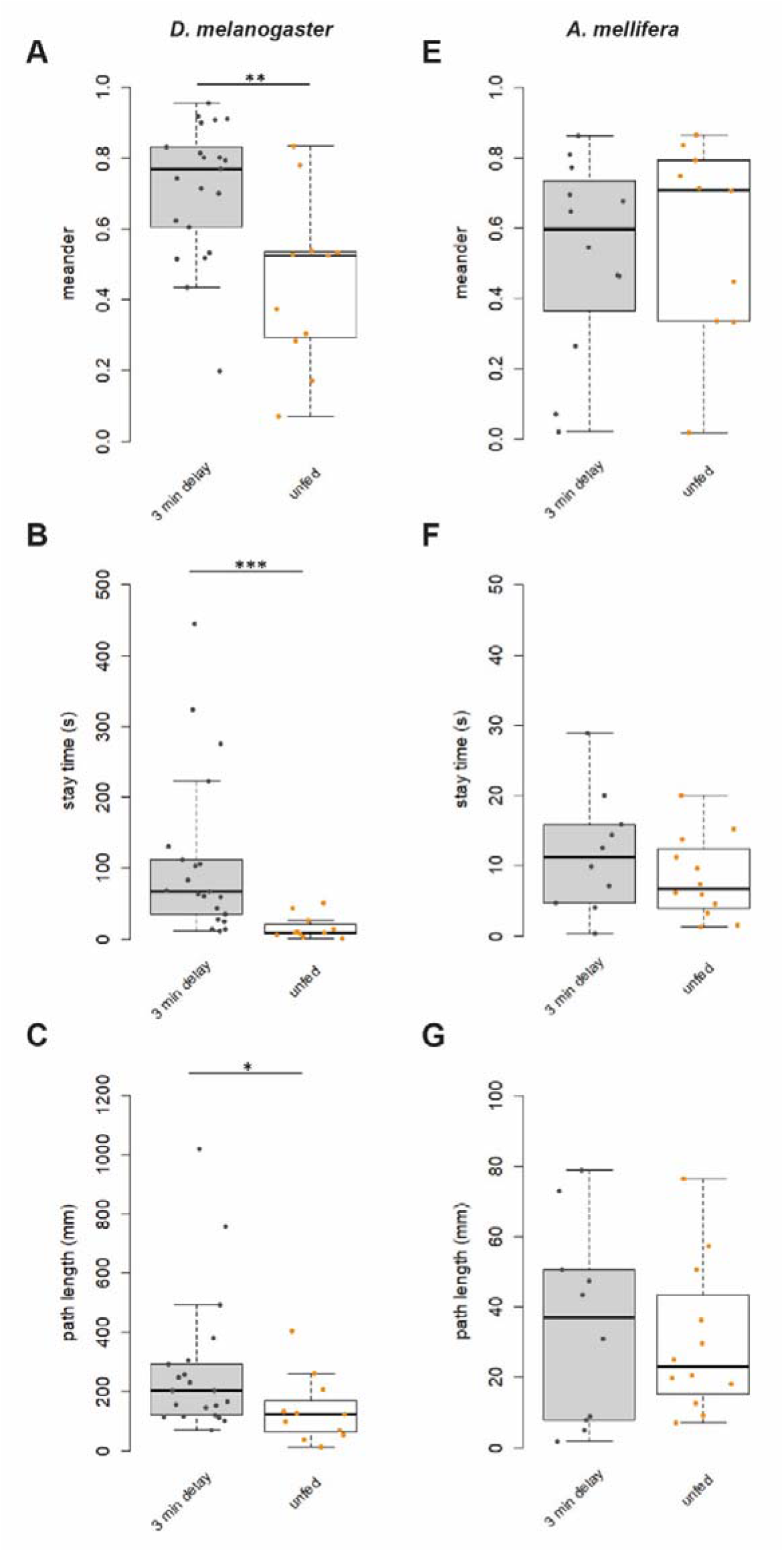
Behavioural parameters of unfed flies and bees compared to 3 min delay; (A-C) Meander, stay time and path length for 3 min delay are significantly higher than unfed controls in flies. (D-F) The parameters did not show a difference between 3 min delay and unfed bees. *p<0.05, **p<0.001, ***p<0.0001, Wilcoxon Rank Sum Test),

**Figure S9:**
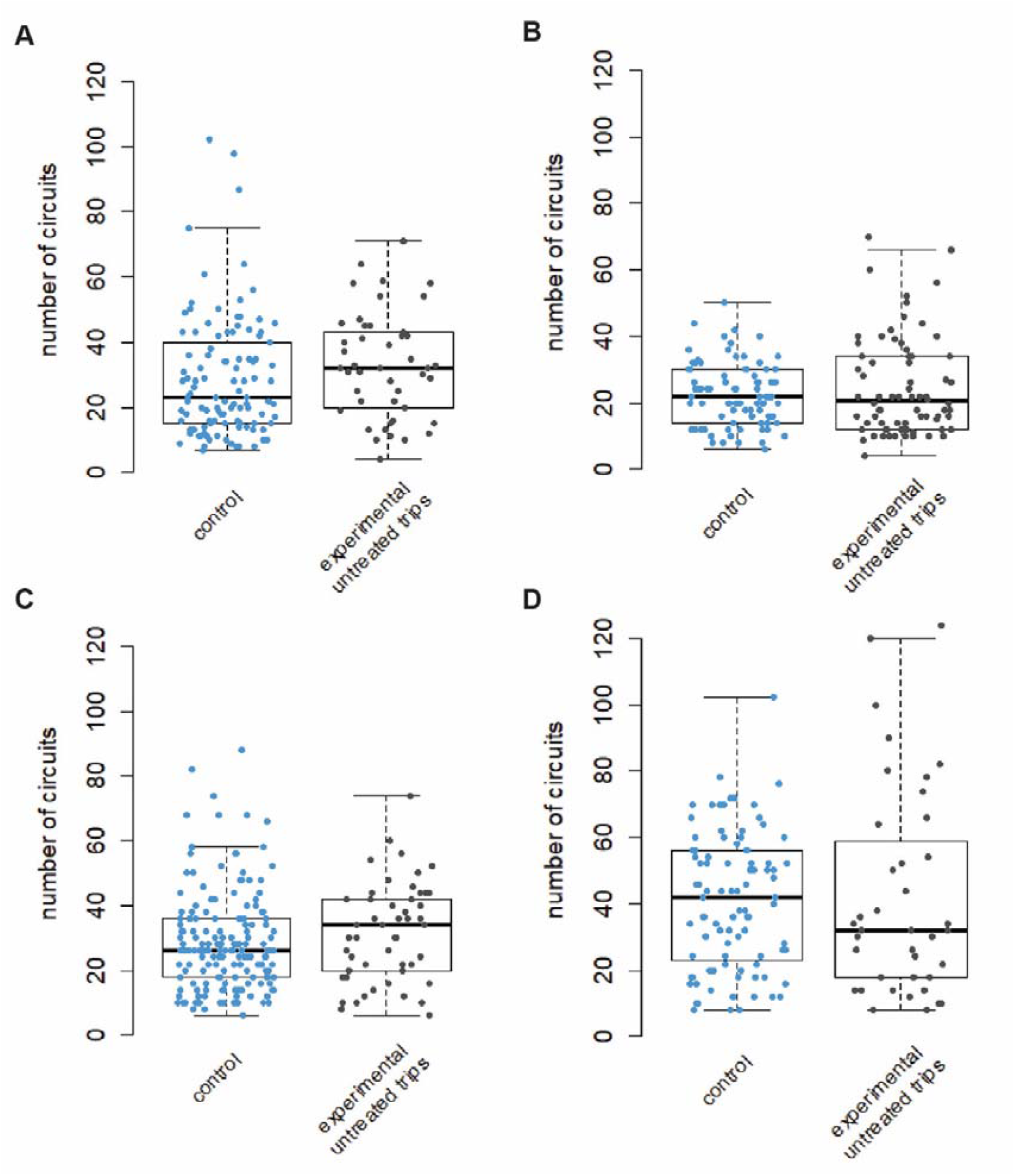
Number of circuits across do not get affected in untreated trips. **(A-D)** The number of circuits for the control group and untreated trips of experimental group (do not show a difference across all four foraging groups (group 1 p=0.07013; group 2 p=0.983; group 3 p=0.1298; group 4 p=0.7505, Wilcoxon Rank Sum Test).

**Figure S10:**
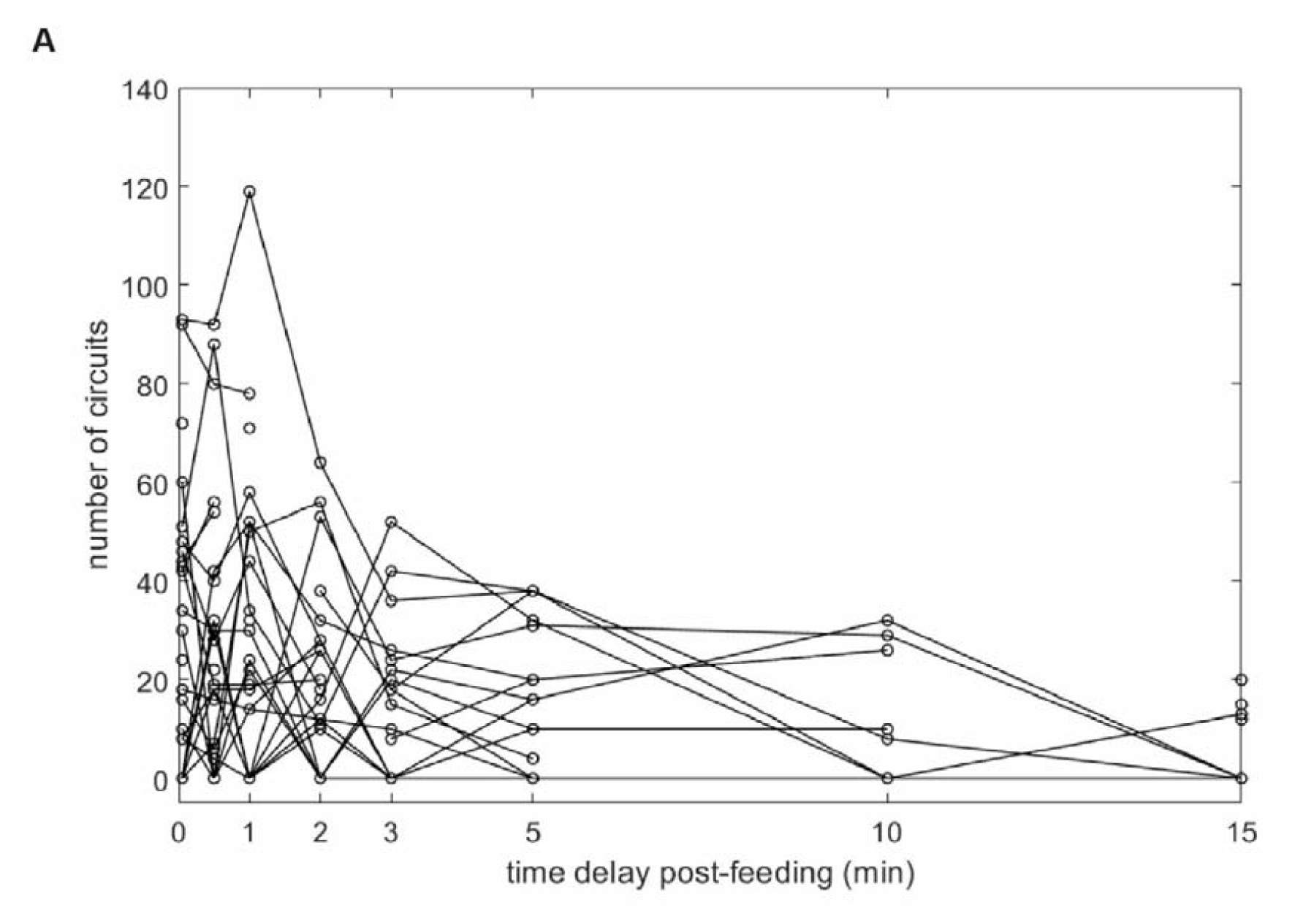
Individual response of each bee for dance experiments. (A) Number of circuits show a general decline with increased delay duration

**Table S1:** Regression analysis on the behavioural parameters were not correlated with the delay durations in flies and bees.

| *D. melanogaster* |  |  |  |  |  |
| --- | --- | --- | --- | --- | --- |
| Parameter | Slope | Intercept | Adjusted R-squared | t-statistic for coefficients | F test p-value |
| number of returns | -0.3204 | 1.4186 | 0.0436 | 1.1172e-10 | 0.0146 |
| meander | -0.05192 | 0.8878 | 0.1634 | 2.2557e-84 | 3.0523e-06 |
| stay time | -66.0444 | 300.03091 | 0.1044 | 1.7434e-19 | 2.6659e-4 |
| path length | -87.5243 | 511.3201 | 0.1122 | 8.2088e-28 | 1.5933e-4 |
| *A. mellifera* |  |  |  |  |  |
| Parameter | Slope | Intercept | Adjusted R-squared | t-statistic for coefficients | F test p-value |
| number of returns | -0.3277 | 1.5798 | 0.0309 | 2.1685e-06 | 0.0931 |
| meander | -0.0749 | 0.7955 | 0.1161 | 3.5176e-28 | 0.0042 |
| stay time | -2.4667 | 21.5779 | 0.0064 | 1.3988e-08 | 0.2434 |
| path length | -11.36 | 72.7391 | 0.0134 | 8.5762e-07 | 0.1833 |

**Table S2:**
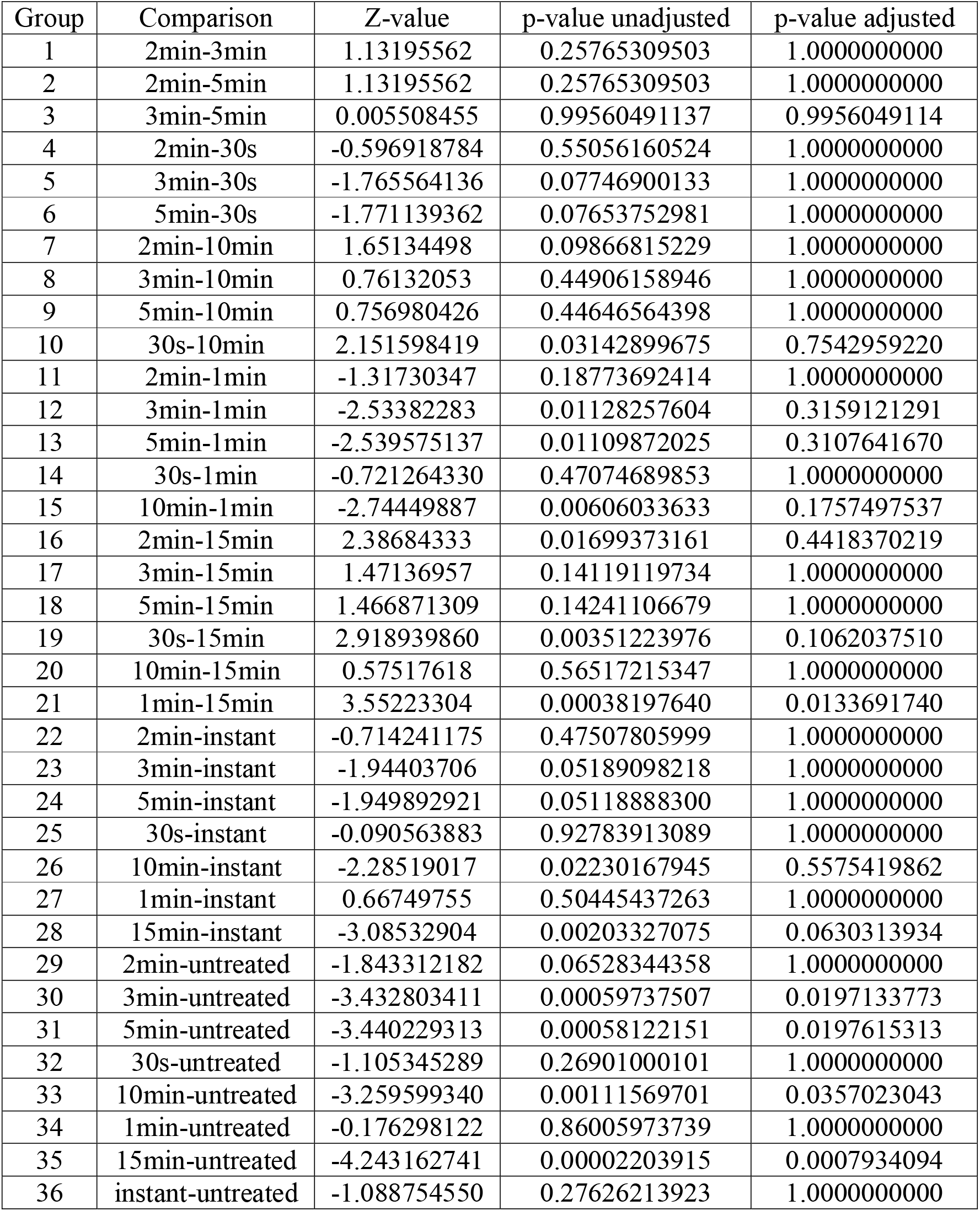
p-values for comparison of number of circuits for dance experiments. Kruskal-Wallis test with Dunn correction, p-values adjusted with the Holm method.

| Group | Comparison | Z-value | p-value unadjusted | p-value adjusted |
| --- | --- | --- | --- | --- |
| 1 | 2min-3min | 1.13195562 | 0.25765309503 | 1.0000000000 |
| 2 | 2min-5min | 1.13195562 | 0.25765309503 | 1.0000000000 |
| 3 | 3min-5min | 0.005508455 | 0.99560491137 | 0.9956049114 |
| 4 | 2min-30s | -0.596918784 | 0.55056160524 | 1.0000000000 |
| 5 | 3min-30s | -1.765564136 | 0.07746900133 | 1.0000000000 |
| 6 | 5min-30s | -1.771139362 | 0.07653752981 | 1.0000000000 |
| 7 | 2min-10min | 1.65134498 | 0.09866815229 | 1.0000000000 |
| 8 | 3min-10min | 0.76132053 | 0.44906158946 | 1.0000000000 |
| 9 | 5min-10min | 0.756980426 | 0.44646564398 | 1.0000000000 |
| 10 | 30s-10min | 2.151598419 | 0.03142899675 | 0.7542959220 |
| 11 | 2min-1min | -1.31730347 | 0.18773692414 | 1.0000000000 |
| 12 | 3min-1min | -2.53382283 | 0.01128257604 | 0.3159121291 |
| 13 | 5min-1min | -2.539575137 | 0.01109872025 | 0.3107641670 |
| 14 | 30s-1min | -0.721264330 | 0.47074689853 | 1.0000000000 |
| 15 | 10min-1min | -2.74449887 | 0.00606033633 | 0.1757497537 |
| 16 | 2min-15min | 2.38684333 | 0.01699373161 | 0.4418370219 |
| 17 | 3min-15min | 1.47136957 | 0.14119119734 | 1.0000000000 |
| 18 | 5min-15min | 1.466871309 | 0.14241106679 | 1.0000000000 |
| 19 | 30s-15min | 2.918939860 | 0.00351223976 | 0.1062037510 |
| 20 | 10min-15min | 0.57517618 | 0.56517215347 | 1.0000000000 |
| 21 | 1min-15min | 3.55223304 | 0.00038197640 | 0.0133691740 |
| 22 | 2min-instant | -0.714241175 | 0.47507805999 | 1.0000000000 |
| 23 | 3min-instant | -1.94403706 | 0.05189098218 | 1.0000000000 |
| 24 | 5min-instant | -1.949892921 | 0.05118888300 | 1.0000000000 |
| 25 | 30s-instant | -0.090563883 | 0.92783913089 | 1.0000000000 |
| 26 | 10min-instant | -2.28519017 | 0.02230167945 | 0.5575419862 |
| 27 | 1min-instant | 0.66749755 | 0.50445437263 | 1.0000000000 |
| 28 | 15min-instant | -3.08532904 | 0.00203327075 | 0.0630313934 |
| 29 | 2min-untreated | -1.843312182 | 0.06528344358 | 1.0000000000 |
| 30 | 3min-untreated | -3.432803411 | 0.00059737507 | 0.0197133773 |
| 31 | 5min-untreated | -3.440229313 | 0.00058122151 | 0.0197615313 |
| 32 | 30s-untreated | -1.105345289 | 0.26901000101 | 1.0000000000 |
| 33 | 10min-untreated | -3.259599340 | 0.00111569701 | 0.0357023043 |
| 34 | 1min-untreated | -0.176298122 | 0.86005973739 | 1.0000000000 |
| 35 | 15min-untreated | -4.243162741 | 0.00002203915 | 0.0007934094 |
| 36 | instant-untreated | -1.088754550 | 0.27626213923 | 1.0000000000 |

## Acknowledgements

The authors are grateful to T. Tanimura for help in developing ideas for the inception of this project. We thank A. Suryanarayan, B. Krishnan, R. Siddaganga, S. Unnikrishnan for their help with the dance experiments. N. Thulasi helped us with the analysis of dance videos. We thank D. Naug, D. Ramesh and S. Unnikrishnan for inputs on earlier versions of the manuscripts.

## Competing interests

The authors declare no competing interests.

## Funding

M.S. was funded by a fellowship by the Indian Council of Medical Research (ICMR). A.B. was supported by National Centre for Biological Sciences, Tata Institute of Fundamental Research institutional funding no. 12P4167.

## Notes

https://data.mendeley.com/datasets/jk26f5tfzk/1

### Competing Interest Statement

The authors have declared no competing interest.

### Summary of Updates

Supplemental files updated

